# Genome-wide analyses of self-reported empathy: correlations with autism, schizophrenia, and anorexia nervosa

**DOI:** 10.1101/050682

**Authors:** Varun Warrier, Roberto Toro, Bhismadev Chakrabarti, the iPSYCH-Broad autism group, Anders D Børglum, Jakob Grove, the 23andMe Research Team, David A. Hinds, Thomas Bourgeron, Simon Baron-Cohen

**Author notes:** **Corresponding authors**: Varun Warrier and Simon Baron-Cohen. Autism Research Centre, Department of Psychiatry, Cambridge University, Douglas House, 18B Trumpington Road, Cambridge, CB2 8AH, United Kingdom. Joint senior authors (David A. Hinds, Thomas Bourgeron, Simon Baron-Cohen).

## Abstract

Empathy is the ability to recognize and respond to the emotional states of other individuals. It is an important psychological process that facilitates navigating social interactions and maintaining relationships, which are important for wellbeing. Several psychological studies have identified difficulties in both self-report and performance-based measures of empathy in a range of psychiatric conditions. To date, no study has systematically investigated the genetic architecture of empathy using genome-wide association studies (GWAS). Here we report the results of the largest GWAS of empathy to date using a well-validated self-report measure of empathy, the Empathy Quotient (EQ), in 46,861 research participants from 23andMe, Inc. We identify 11 suggestive loci (P < 1×10^-6^), though none were significant at P < 2.5×10^-8^ after correcting for multiple testing. The most significant SNP was identified in the non-stratified analysis (rs4882760; P = 4.29×10^-8^), and is an intronic SNP in *TMEM132C.* The EQ had a modest but significant narrow-sense heritability (0.11±0.014; P = 1.7×10^-14^). As predicted, based on earlier work, we confirmed a significant female-advantage on the EQ (P < 2×10^-16^ Cohen’s d = 0.65). We identified similar SNP heritability and high genetic correlation between the sexes. Also, as predicted, we identified a significant negative genetic correlation between autism and the EQ (r_g_ = -0.27±0.07, P = 1.63×10^-4^). We also identified a significant positive genetic correlation between the EQ and risk for schizophrenia (r_g_ = 0.19±0.04; P= 1.36×10^-5^), risk for anorexia nervosa (r_g_ = 0.32±0.09; P = 6×10^-4^), and extraversion (r_g_ = 0.45±0.08; 5.7×10^-8^). This is the first GWAS of self-reported empathy. The results suggest that the genetic variations associated with empathy also play a role in psychiatric conditions and psychological traits.

## Introduction

Empathy is the ability to identify other people’s thoughts, intentions, desires, and feelings, and to respond to others’ mental states with an appropriate emotion^1^. It plays an important role in social interaction by facilitating both making sense of other people’s behaviour and in responding appropriately to their behaviour. For these reasons, it is considered a key component of prosocial behaviour, social cooperation, and social cognition^2^. Aspects of empathy are observed in humans and other animals and is thought to have evolved to support a range of prosocial behaviour and cooperative behaviour^2^.

Differences in various fractions empathy have been observed in several psychiatric conditions including autism^1^, bipolar disorder^3^, schizophrenia^4–6^, and major depressive disorder^3,7,8^. Two major fractions of empathy include affective empathy (the drive to respond to another’s mental state with an appropriate emotion) and cognitive empathy (the ability to recognize another’s mental state). These differences vary between psychiatric conditions: for example, individuals with schizophrenia are more likely to report higher personal distress and emotional contagion^6^, whereas individuals with autism are likely to show difficulties with cognitive empathy but not affective empathy^1,9^. These may reflect a causal risk mechanisms where alterations in empathy contribute to higher risk for developing a psychiatric condition. Equally, differences in empathy may also be due to the presence of a psychiatric condition, which may not allow individuals to understand and respond to another person’s mental state effectively.

Whilst empathy is clearly shaped by early experience, parenting, and other social factors, different lines of evidence suggest that empathy is partly biological. Empathy is modestly heritable (approximately a third of the variance is heritable)^10–12^, and a few candidate gene association studies have investigated the role of various genes in empathy^13-15^. In addition, several studies have identified a role for the oxytocinergic and the foetal testosterone systems in modulating empathy^2,16–18^. Neuroimaging studies have identified distinct brain regions implicated in different aspects of empathy including the amygdala and the ventromedial prefrontal cortex^19,20^. Empathy also shows a marked sex difference: females, on average, score higher on different measures of empathy^1,21^. A longitudinal study suggests that this female advantage grows larger with age^22^. Sex differences in the mind arise from a combination of innate biological differences, cultural, and environmental differences. Studies in infant humans have identified sex differences in the developmental precursors to empathy, such as neonatal preferences to faces over objects^23^ when environmental and cultural influences are minimal, lending support to the idea that sex differences in empathy are at least partly biological^24^.

Because empathy difficulties are found in a range of psychiatric conditions, it is an important phenotype for investigation. Understanding the biological networks that partly determine empathy may help us understand how it contributes to psychiatric phenotypes, an approach that has been used for other traits such as neuroticism^25^, creativity^26^, and cognitive ability^27^. We investigate the genetic architecture of self-reported empathy using the Empathy Quotient (EQ)^1^. The EQ is listed in the Research Domain Criteria (RDoC)^28^ as a self-report measure under the domain of ‘Understanding Mental States’ (https://www.nimh.nih.gov/research-priorities/rdoc/units/self-reports/151133.shtml). The EQ is widely used, has excellent test-retest reliability (r = 0.83, P = 0.0001)^29^, high internal consistency (Cronbach’s alpha ~ 0.9)^30,31^ and is significantly correlated with factors in the Interpersonal Reactivity Index and the Toronto Empathy Questionnaire, two other measures of empathy, suggesting good concurrent validity^29,32^. Psychometric analysis of the EQ in 3,334 individuals from the general population suggests that the EQ is a good measure of empathy which can be measured across a single dimension^33^. By focusing on a self-report measure of empathy, we were able to obtain phenotypic and genetic data from a large number of participants, increasing the statistical power of the study. Previous work from our lab investigated the genetic correlates of cognitive empathy using a specific performance measure – the ‘Reading the Mind in the Eyes’ Test (the Eyes Test)^34^. The Eyes Test has a low correlation with the EQ (r ~ 0.10)^30,35^ and measures only one facet of empathy, namely cognitive empathy. Cognitive empathy is also referred to as employing a ‘theory of mind’, or ‘mentalizing’. The EQ includes items that measure cognitive empathy, but others that measure affective empathy. Yet other items on the EQ involve both cognitive and affective empathy.

In this study, we aim to answer three questions: 1. What is the genetic architecture of empathy? 2. Is empathy genetically correlated to various psychiatric conditions, psychological traits, and education? and 3. Is there a genetic contribution to sex differences in empathy? We performed sex-stratified and non-stratified genome-wide association analyses of empathy in research participants from 23andMe, a personalized genetics company. We calculated the narrow sense heritability explained by all the SNPs tested, and investigated sex differences. Finally, we conducted genetic correlation analyses with six psychiatric conditions (anorexia, ADHD, autism, bipolar disorder, major depressive disorder, and schizophrenia), psychological traits, and educational attainment.

## Methods

### Participants

Research Participants were drawn from the customer base of 23andMe, Inc. a personal genetics company and are described in detail elsewhere^36,37^. There were 46,861 participants (24,543 females and 22,318 males). All participants included in the analyses provided informed consent and answered surveys online according to a human subjects research protocol, which was reviewed and approved by Ethical & Independent Review Services, an AAHRPP-accredited private institutional review board (http://www.eandireview.com). All participants completed the online version of the questionnaire accessible via the research tab of their password protected 23andMe personal online account. Only participants who were primarily of European ancestry (97% European Ancestry) were selected for the analysis using existing methods^38^. Unrelated individuals were selected using a segmental identity-by-descent algorithm^39^.

### Measures

The Empathy Quotient (EQ)^1^, is a self-report measure of empathy, and includes items relevant to both cognitive and affective empathy. It comprises 60 questions and has a good test-retest reliability^29^. 20 questions are filler questions, of the remaining 40 questions participants can score a maximum of 2 points and a minimum of 0 point per question. Therefore, in this study, participants scored a maximum of 80 and a minimum of 0.

### Genotyping, imputation and quality control

DNA extraction and genotyping were performed on saliva samples by the National Genetic Institute, USA. Participants were genotyped on one of four different platforms – V1, V2, V3 and V4. The V1 and V2 platforms have a total of 560,000 SNPs largely based on the Illumina HumanHap550+ BeadChip. The V3 platform has 950,000 SNPs based on the Illumina OmniExpress+ Beadchip and has custom content to improve the overlap with the V2 platform. The V4 platform is a fully customized array and has about 570,000 SNPs. All samples had a call rate greater than 98.5%. A total of 1,030,430 SNPs (including Insertion/Deletion or InDels) were genotyped across all platforms. Imputation was performed using the March 2012 (v3) release of the 1000 Genomes Phase 1 reference haplotypes. First, we used Beagle (version 3.3.1)^40^ to phase batches of 8000-9000 individuals across chromosomal segments of no more than 10,000 genotyped SNPs, with overlaps of 200 SNPs. SNPs were excluded if they were not in Hardy-Weinberg equilibrium (P < 10^-20^), had a genotype call rate less than 95%, or had discrepancies in allele frequency compared to the reference European 1000 Genomes data (chi-squared P < 10^-15^). We then imputed each phased segment against all-ethnicity 1000 Genomes haplotypes (excluding monomorphic and singleton sites) using Minimac2^41^, using 5 rounds and 200 states for parameter estimation. We restricted the analyses to only SNPs that had a minor allele frequency (MAF) of at least 1%. For genotyped SNPs, those present only on platform V1, or in chromosome Y and mitochondrial chromosomes were excluded due to small sample sizes and unreliable genotype calling respectively. Next, using trio data from all research participants in the 23andMe dataset, where available, SNPs that failed a parent offspring transmission test were excluded. For imputed SNPs, we excluded SNPs with average r^2^ < 0.5 or minimum r^2^ < 0.3 in any imputation batch, as well as SNPs that had strong evidence of an imputation batch effect. The batch effect test is an F test from an ANOVA of the SNP dosages against a factor representing imputation batch; we excluded results with P < 10^−50^. After quality control, 9,955,952 SNPs were analysed. Genotyping, imputation, and preliminary quality control were performed by 23andMe.

### Genetic association

We performed a linear regression assuming an additive model of genetic effects. Age and sex along with the first five ancestry principal components were included as covariates. Additionally, we performed a male-only and a female-only linear regression analysis to identify sex-specific loci. Since we were performing two independent tests for each trait (male-only and female-only, and males and females combined with sex as a covariate which is equivalent to a meta-analysis of the two sex-stratified GWAS), we used a threshold of P < 2.5×10^-8^ (5×10^-8^/2) to identify significant SNPs. Leading SNPs in each locus was identified after pruning for LD (r^2^ > 0.8) using Plink version 1.9. We calculated the variance explained by the top SNPs using a previously used formula^42^:

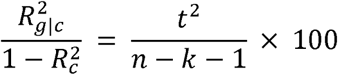

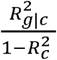 is the proportion of variance explained by the SNP after accounting for the effects of the covariates (four ancestry principal components, age, and, additionally, sex for the non-stratified analyses), t is the t-statistic of the regression co-efficient, k is the number of covariates, and n is the sample size. Winner’s curse correction was conducted using FDR Inverse Quantile Transformation^43^.

### Genomic inflation factor, heritability, and functional enrichment

We used Linkage Disequilibrium Score regression coefficient (LDSR) to calculate genomic inflation due to population stratification^44^ (https://github.com/bulik/ldsc). Heritability and genetic correlation was performed using extended methods in LDSR^45^. Difference in heritability between males and females was quantified using^46^:

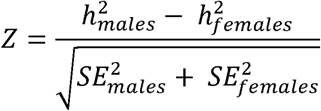

where Z is the Z score for the difference in heritability for a trait, (h^2^_males_ - h^2^_females_) is the difference SNP heritability estimate in males and females, and SE is the standard errors for heritability. Two-tailed P-values were calculated, and reported as significant if P < 0.05. This We identified enrichment in genomic functional elements for the traits by partitioning heritability performed in LDSR^47^. In addition to the baseline partitions we conducted four additional enrichment analyses. Enrichment for CNS specific histone marks was conducted using cell type specific partitioned heritability analysis. For genes that are intolerant to loss-of-function mutations, we identified gene boundaries of genes with probability of loss-of-function intolerance scores > 0.9 from the Exome Aggregation Consortium^48^, and conducted partitioned heritability analysis for all common SNPs within the gene boundaries identified. Similarly, for sex-differentially enriched genes, we identified gene boundaries of genes with sex differential expression in cerebral cortex and associates structures (Brain Other)^49^ **(Supplementary Table 8)**. We divided this into two separate lists – genes with higher expression in males and genes with higher expression in females with an FDR corrected P-value < 0.05. Partitioned heritability analyses were conducted to identify enrichment using LDSR.

### Genetic correlations

LDSR was also used to calculate genetic correlations. We restricted our analyses to only the non-stratified GWAS dataset due to the unavailability of sex-stratified GWAS data in the phenotypes investigated. We calculated initial genetic correlations using LD Hub^50^ for schizophrenia^51^, bipolar disorder, major depressive disorder, depressive symptoms, educational attainment (years of schooling.), NEO-Openness to experience, NEO-Conscientiousness, subjective wellbeing, and neuroticism. For anorexia nervosa^52^, and ADHD^53^, we used the data available from the PGC webpage (https://www.med.unc.edu/pgc/results-and-downloads) to conduct genetic correlation analyses as these are in larger samples and, consequently, have greater statistical power than the datasets available on LD Hub. For autism, we used summary statistics from the PGC-iPSYCH meta-analysis^54^, details of which are provided in the **Supplementary Note**. We also conducted genetic correlation for extraversion^55^ separately as the data was unavailable on LD Hub. For the anorexia nervosa, autism and the extraversion analyses, the North West European LD scores were used and the intercepts were not constrained as the extent of participant overlap was unknown. We report significant lists if they Bonferroni corrected P < 0.05, which we acknowledge is conservative. For anorexia nervosa and autism, we also conducted genetic correlation analyses using the sex-stratified EQ dataset due to the significant sex-differences observed in these conditions. We correct for these using Bonferroni correction, and report significant correlations at P < 0.05.

### Gene-based analysis

Gene based analyses for the non-stratified GWAS were performed using MAGMA^56^, which integrates LD information between SNPs to prioritize genes. Genes were significant if they had a Bonferroni corrected P < 0.05. In addition, we also investigated enrichment in Gene Ontology terms using MAGMA.

### Genome-wide colocalization

Pairwise genome-wide colocalization analyses were conducted using GWAS-PW^57^ by dividing the genomes into segments containing approximately 5,000 SNPs each. We considered the posterior probability of model 3 i.e. the model wherein SNPs in the same locus influence both the traits. We used a rigorous threshold of posterior probability > 0.95 to identify significant loci that influenced both the traits. We conducted pairwise colocalization for empathy (non-stratified), and schizophrenia^51^, anorexia nervosa^52^, and autism.

### Data Availability

Summary statistics for the EQ GWAS can be requested directly from 23andMe, and will be made available to qualified researchers subject to the terms of a data transfer agreement with 23andMe that protects the privacy of the 23andMe research participants. Please contact David Hinds for more information. Top SNPs can be visualized here: https://ghfc.pasteur.fr/eq/.

## Results

### Phenotype description

To understand the genetic architecture of empathy, we collaborated with 23andMe to conduct a Genome Wide Association Study (GWAS) of empathy (n = 46,861) using the EQ, which was normally distributed. A flow chart of the study protocol is shown in **Figure 1**. The mean score for all participants was 46.4 (sd = 13.7) on a total of 80 on the EQ, which is similar to the mean score reported in 90 typical participants in the first study describing the EQ (42.1, sd =10.6)^1^. The mean age of the participants was 48.9 (sd = 15.7). Females scored higher than males on the EQ (41.9±13.5 in males, 50.4±12.6 in females) **(Figure 2)**, as previously observed^1^. There was significant age effect, with scores increasing with age (Beta = 0.08±0.003; P = 3.3×10^-104^) and a significant sex effect with females scoring higher than males (Beta = 8.4±0.11; P ~ 0).

**Figure 1.**
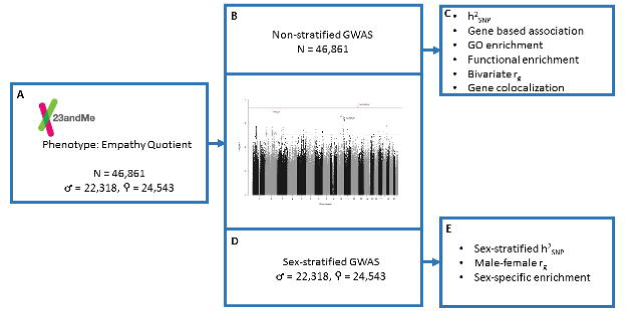
Schematic diagram of the study protocol. Phenotyping was conducted in research participants from 23andMe., Inc using the 60 question Empathy Quotient (Panel A). Three GWAS analyses were conducted: non-stratified GWAS (Panel B) and sex-stratified GWAS (Panel D) in unrelated individuals of primarily European ancestry. Summary non-stratified GWAS data was used to conduct SNP heritability, gene, pathway, and functional enrichment, genetic correlations with psychiatric conditions, psychological traits and education, and Bayesian gene colocalization (Panel C). Summary sex-stratified GWAS data was used to conduct sex-specific SNP heritability, genetic correlation between the male and female datasets, and enrichment in sex-differentially expressed genes (Panel E).

### Genome-wide association analyses

We conducted three GWAS analyses: a male-only analysis, a female-only analysis, and a non-stratified analysis, using a linear regression model with age and the first four ancestry principal components as covariates **(Methods)**. We corrected for the three different tests using a conservative threshold of P = 2.5×10^-8^. LD score regression coefficient suggested non-significant genomic inflation due to population stratification **(Supplementary Figure 1 – 3)**. We did not identify any genome wide significant SNPs **(Supplementary Figures 1 - 3 and Supplementary Table 1)**. We identified 11 suggestive loci (P < 1 x 10^-6^) in the three GWAS. The most significant SNP was identified in the non-stratified analysis (rs4882760; P = 4.29×10^-8^), and is an intronic SNP in *TMEM132C*. Regional association plots for all suggestive loci are provided in **Supplementary Figure 4**.

To investigate if the top SNPs from the EQ also contribute to cognitive empathy as measured by the Eyes Test, we conducted SNP lookup of all the 11 suggestive loci in the Eyes Test GWAS. None of the 11 loci were significant in the Eyes Test GWAS, and only 7 out of the 11 SNPs had concordant effect directions in the two traits (P = 0.54; two-sided binomial sign test).

The most significant SNP in each GWAS analysis explained 0.06 – 0.13% of the total variance **(Supplementary Table 2).** However, this reduced to 0.0006 – 0.016% after correcting for winner’s curse **(Supplementary Table 2)**.

### Gene-based association, heritability, and enrichment in functional categories

Gene based analysis identified two significant genes for the EQ: *SEMA6D* (P = 9.14×10^-7^), and *FBN2* (P = 1.68×10^-6^) **(Supplementary Table 3)**. Analysis for enrichment in Gene Ontology (GO) terms did not identify any significant enrichment **(Supplementary Table 4)**. The most significant GO process was negative regulation of neurotransmitter secretion.

We used LDSR^44^ to calculate the heritability explained by all the SNPs tested (Methods) and identified a heritability of 0.11±0.014 for the EQ (P = 1.7×10^-14^) **(Figure 2, Supplementary Table 5)**. Partitioning heritability by functional categories did not identify any significant enrichment after correcting for multiple testing **(Supplementary Table 6)**. We also investigated if there was an enrichment in heritability for histone marks in cells in the CNS^47^, but did not find a significant enrichment (enrichment = 3.67±1.45; P = 0.077).

Recent studies have identified an enrichment of associations in or near genes that are extremely intolerant to loss-of-function variation in schizophrenia^58^, autism^59,60^, and developmental disorders^61^, conditions that are often accompanied by difficulties in social behaviour and empathy. We investigated if there was a significant enrichment in GWAS signal for the EQ in these genes that are extremely intolerant to loss-of-function variation. We did not identify a significant enrichment after correction for multiple testing (proportion h^2^_SNP_ =0.19, proportion SNP = 0.09, enrichment = 1.83±0.42; P = 0.044).

**Figure 2.**
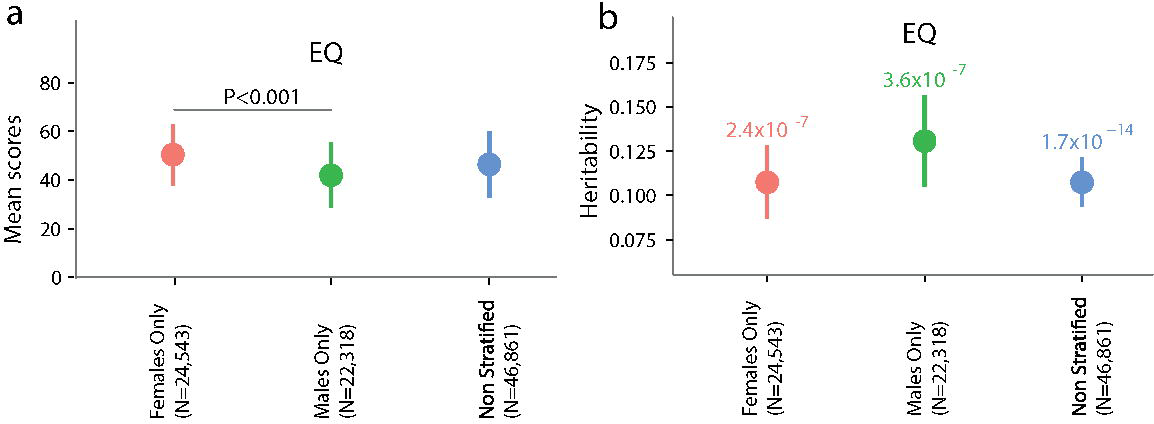
Mean scores and heritability estimates for the EQ. Mean scores and standard deviations for scores on the EQ (a). The effect size of the difference between males and females for the EQ scores was Cohen’s d= 0.65. The bar on top and the number represents the statistical significance of the male-female difference in mean scores for the EQ. Mean estimates and standard errors for heritability for scores on the EQ (b) for the females-only GWAS, males-only GWAS and the non-stratified GWAS. Numbers on top of the graphs represent P-values for each heritability estimate. Note, the y-axis does not start at 0.

### Sex differences

Sex differences in empathy^62^ may reflect genetic as well as non-genetic factors (such as prenatal steroid hormones, and postnatal learning)^63^. In our dataset, there was a significant female advantage on the EQ (P < 2×10^-16^ Cohen’s d = 0.65) **(Figure 2)**. To investigate the biological basis for the sex-difference observed in the traits, we first tested the heritability of the sex-stratified GWAS analyses for the EQ. Our analyses revealed no significance difference between the heritability in the males-only and the females-only datasets (P = 0.48 for male-female difference in the EQ) **(Figure 2 and Supplementary Table 5)**. Additionally, there was a high genetic-correlation between the males-only and females-only GWAS (r_g_ = 0.82±0.16, P = 2.34×10^-7^), indicating a high degree of similarity in the genetic architecture of the traits in males and females. This was not significantly different from 1 (P = 0.13, one-sided Wald Test). We investigated the heterogeneity in the 11 SNPs of suggestive significance in both the sexes using Cochran’s Q-Test, and did not identify significant heterogeneity **(Supplementary Table 7)**.

Sex differences may also arise by differential expression of specific genes in different neural tissues at different developmental stages^49,64^. This could be due to multiple factors, including sex-specific transcription factors and sex-specific DNA methylation. We investigated this by performing enrichment analysis of the non-stratified GWAS in genes with higher expression in either males or females in cortical tissue samples **(Supplementary Table 8)**. We did not identify a significant enrichment for either genes with higher expression in males (enrichment = 1.95±0.70, P = 0.17) or females (enrichment = 0.28±0.84, P = 0.39).

### Genetic correlations

To investigate how the EQ correlates with psychiatric conditions, psychological traits and educational attainment, we performed genetic correlation **(Methods)** with six psychiatric conditions (autism, ADHD, anorexia nervosa, bipolar disorder, depression (major depressive disorder and the larger depressive symptoms dataset) and schizophrenia), six psychological traits (NEO-extraversion, NEO-openness to experience, NEO-conscientiousness, neuroticism, and subjective wellbeing), and educational attainment (a proxy measure of IQ, measured using years of schooling) **(Supplementary Table 9)**. With psychiatric conditions, three genetic correlations were significant following Bonferroni correction: EQ-autism (r_g_ = -0.27 ± 0.07, P = 1.63×10^-4^), EQ-schizophrenia (r_g_ = 0.19±0.04; P = 1.36×10^-5^) and EQ-anorexia nervosa (r_g_ = 0.32±0.09; P = 6×10^-4^) (**Figure 3**).

As anorexia nervosa is primarily diagnosed in women, and autism is primarily diagnosed in men, we further tested sex-specific correlations. After Bonferroni correction, we identified significant genetic correlations between the EQ in females (EQ-F) and anorexia (r_g_ = 0.48±0.12; P = 8.46×10^-5^) and the EQ in males (EQ-M) and autism (r_g_ = -0.3±0.08, P = 3×10^-4^).

With psychological traits, we identified one significant correlation after Bonferroni correction: EQ with extraversion (r_g_ = 0.45±0.08; P = 5.76×10^-8^). Additionally, we identified two nominally significant correlations: the EQ with subjective wellbeing (r_g_ = 0.19±0.07; P = 7.8×10^-3^), and NEO-conscientiousness (r_g_ = 0.39±0.14; P = 8.8×10^-03^) (**Figure 3**). All three correlations were in the predicted direction as studies have identified a positive phenotypic correlation between all three traits and the EQ^31,65^. We previously reported a small positive correlation between the EQ and the Eyes Test (r_g_ = 0.18±0.06; P = 0.007)^34^, mirroring previous reported estimates of phenotypic correlation in the general population^35^ and estimates in our database from 916 neurotypical adults (r = 0.11±0.032; P = 0.003, Pearson correlation).

### Bayesian genomic colocalization

As there were significant genetic correlations between the EQ, anorexia nervosa schizophrenia, and autism, we investigated if there are genomic regions that influence both empathy and one of the psychiatric conditions (colocalization) by estimating the Bayesian posterior probability. We did not identify any regions associated with empathy and the three conditions. The most significant region identified in this analysis was in Chr11p12, posterior probability = 0.78 (**Supplementary Figure 5)** in the empathy-anorexia analysis. The most significant SNPs in this region for both anorexia and empathy are intronic SNPs in the gene *LRRC4C*, which is implicated in excitatory synapse development^66,67^. Further, this gene is highly intolerant to loss-of-function mutations (probability of Loss-of-function Intolerance = 0.95). We did not identify any eQTLs in this region in neural tissues. These results are preliminary, and a cautious interpretation is warranted as the probability is influenced by the modest power of both the GWAS^57^.

**Figure 3.**
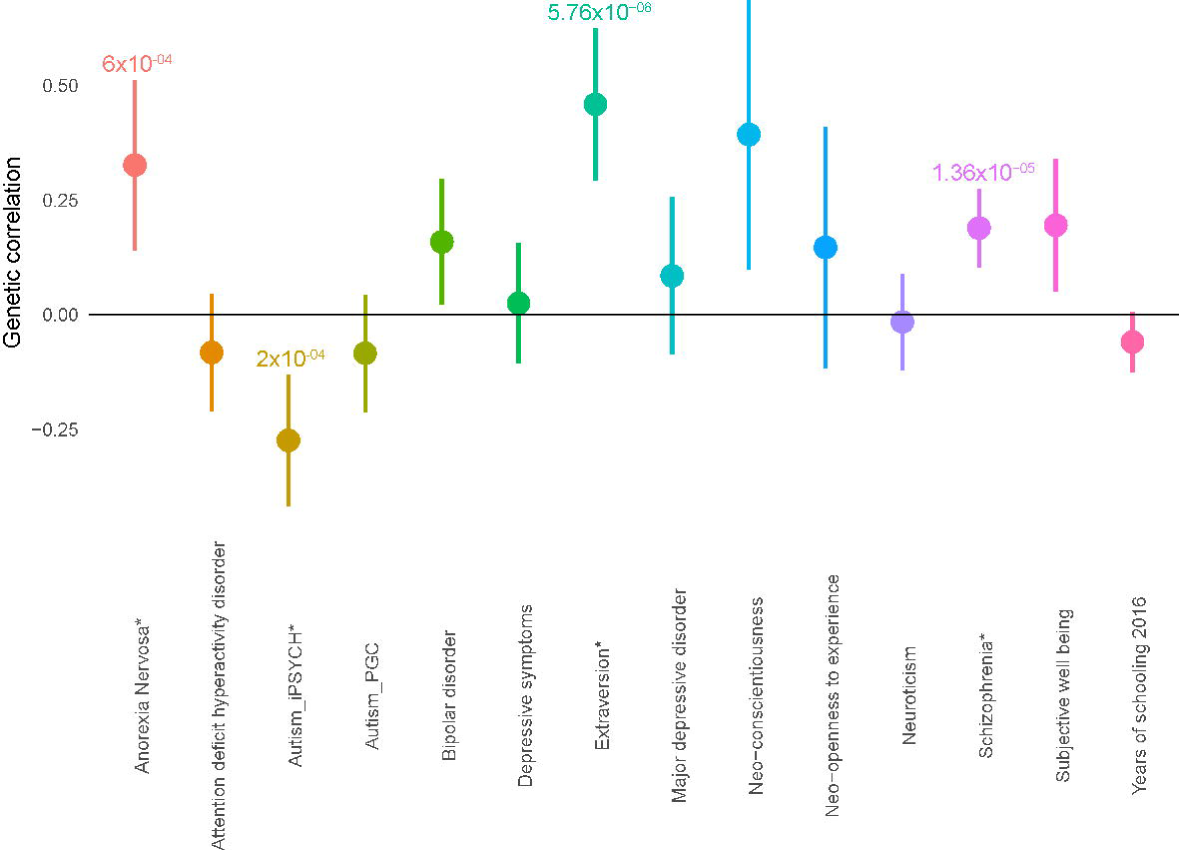
Genetic correlations between the EQ and other conditions. Mean and 95% confidence intervals shown for genetic correlations between empathy and other conditions. P-values are provided significant correlations after Bonferroni correction.

## Discussion

This is the first GWAS to investigate the genetic architecture of self-reported empathy. We identified three significant genetic correlations with the EQ and psychiatric conditions and psychological traits (anorexia nervosa, schizophrenia, and extraversion), providing insights into the shared genetic architecture. Although we did not identify any significant SNPs after correcting for multiple testing at P < 2.5×10^-8^, we identified eleven SNPs of suggestive significance (P < 1×10^-6^_)_. Males and females perform differently on the tests, but there was limited evidence of a sex-specific genetic architecture.

We identified a significant negative genetic correlation between the EQ and autism. Several studies have identified lower self-reported empathy in individuals with autism, and our results mirror these studies^1,68^. This is likely to be driven by difficulties in understanding the mental states of others rather than responding to them. We also identified significant genetic correlations for the EQ with schizophrenia and anorexia nervosa. The empirical literature in general report deficits in cognitive empathy^4,70^, but preserved or stronger affective empathy^6,70^ and emotional contagion/personal distress^6^ in individuals with schizophrenia compared to controls. Studies with anorexia nervosa, on the other hand, have yielded mixed results. Some studies suggest preserved empathy^71^, some identify reduced cognitive empathy^72–74^, and others identify greater emotional contagion/personal distress^75^ in individuals with anorexia nervosa compared to controls. These studies are typically conducted in small samples, which may explain the different results in these heterogeneous conditions. Our results suggest that genetic variants associated with self-reported empathy slightly increase the risk for schizophrenia and anorexia nervosa; the latter remained significant even after using the females-only EQ dataset. A previous study^34^ identified a significant genetic correlation between cognitive empathy (measured using the Eyes Test^76^) and anorexia nervosa, underscoring the importance of empathy as a genetic risk factor in anorexia nervosa. However, both cognitive empathy and anorexia nervosa are positively correlated with educational attainment^34,77^, and it is possible that the correlation between cognitive empathy and anorexia nervosa may be mediated by educational attainment.

Here, self-reported empathy is not genetically correlated with educational attainment, which allows us to conclude that empathy indeed contributes to genetic risk for anorexia nervosa. Further, while cognitive empathy was not correlated with schizophrenia, self-reported empathy was positively and significantly correlated with schizophrenia, suggesting distinct roles for the two phenotypes in pathology. Schizophrenia and anorexia share significant positive genetic correlation (r_g_ = 0.23±0.06)^77^, and it is possible that the pleiotropy between these two conditions may, in part, be mediated by genetic variants that contribute to empathy. This needs to be tested. The modest power of the remaining psychiatric GWAS precludes the identification of significant genetic correlations with self-reported empathy. Together with the GWAS on cognitive empathy^34^, this study provides evidence for the distinct roles of different social processes in various psychiatric conditions.

Investigating genetic correlation results with psychological traits and measures of cognition further helped elucidate the genetic architecture of self-reported empathy. The EQ was significantly correlated with extraversion and nominally correlated with subjective wellbeing and conscientiousness. Both extraversion and conscientiousness correlate with empathy^65^ which, in turn, contributes to subjective wellbeing^31^. Of the five personality factors, extraversion, conscientiousness and agreeableness have modest correlations with self-reported empathy^65^. We did not test for genetic correlation with agreeableness due to the low heritability of the trait. The direction of our genetic correlation results mirror observed phenotypic correlations and provide additional evidence for the positive role of self-reported empathy in subjective wellbeing.

This is also the first study to provide estimates of additive heritability explained by all the SNPs tested for self-reported empathy, and approximately 11% of variance was explained by SNPs. One study, investigating the heritability of the reduced EQ (18 items) in 250 twin pairs, identified a heritability of 0.32^12^. The literature on the heritability of empathy and prosociality is inconsistent, with heritability estimates ranging from 0.20^10^ to 0.69^78^, although a meta-analysis of different studies identified a heritability estimate of 0.35 (95% CI – 0.21 – 0.41)^79^. Our analysis therefore suggests that a third of the heritability can be attributed to common genetic variants. Like IQ^80^, the heritability of empathy and prosociality behaviour changes with age^10^. We did not investigate the effect of age on heritability in our study.

We did not find any significant differences in heritability between males and females. Further, the male-female genetic correlations for both the traits were high. Despite the high genetic correlation, sex-specific correlations with anorexia nervosa were significant only for the females-only dataset. This suggests that the sex-specific genetic component of empathy can contribute differentially to psychiatric conditions. Several other factors may explain the observed phenotypic sex difference. For example, genetic variants for empathy may be enriched in sex-specific gene expression pathways. We conducted preliminary analysis by investigating if there is an enrichment in sex-differentially expressed genes in cortical tissue samples, but did not find significant enrichment. However, sex-specific gene expression is a dynamic process with both spatial and developmental differences^49,64^. Investigating across different tissues and developmental time points in well powered gene expression datasets will help better understand sex differences in empathy.

There are a few limitations that need to be taken into consideration in interpreting our results. The EQ is a self-report measure, and while it has excellent psychometric properties and construct validity, and it is unclear how much of the intrinsic biological variation in this trait is captured by it. Further, while this is the largest GWAS to date of self-reported empathy, it still is only modestly powered, reflected in our inability to identify genome-wide significant loci. This modest statistical power influences subsequent analysis, and we highlight several nominally significant results for further investigation in larger datasets.

In conclusion, the current study provides the first narrow sense heritability for empathy. While there is a highly significant difference on the EQ between males and females, heritability is similar, with a high genetic correlation between the sexes. We also identified significant genetic correlations between empathy and some psychiatric conditions and psychological traits, including autism. This global view of the genomic architecture of empathy will allow us to better understand psychiatric conditions, and improve our knowledge of the biological bases of neurodiversity in humans.

## Acknowledgements

We thank Richard Bethlehem, Florina Uzefovsky, and Paula Smith for discussions of the results. We are grateful to Brendan Bulik-Sullivan, Hillary Finucane, and Donna Werling for their help with the analytical methods. This study was funded by grants from the Medical Research Council, the Wellcome Trust, the Autism Research Trust, the Templeton World Charity Foundation, the Institut Pasteur, the CNRS and the University Paris Diderot. VW is funded by St. John’s College, Cambridge, and Cambridge Commonwealth Trust. The research was funded and supported by the National Institute for Health Research (NIHR) Collaboration for Leadership in Applied Health Research and Care East of England at Cambridgeshire and Peterborough NHS Foundation Trust. The views expressed are those of the author(s) and not necessarily those of the NHS, the NIHR or the Department of Health. We would like to thank the research participants and employees of 23andMe for making this work possible. We specifically thank the following members of the 23andMe Research Team: Michelle Agee, Babak Alipanahi, Adam Auton, Robert K. Bell, Katarzyna Bryc, Sarah L. Elson, Pierre Fontanillas, Nicholas A. Furlotte, Bethann S. Hromatka, Karen E. Huber, Aaron Kleinman, Nadia K. Litterman, Matthew H. McIntyre, Joanna L. Mountain, Carrie A.M. Northover, Steven J. Pitts, J. Fah Sathirapongsasuti, Olga V. Sazonova, Janie F. Shelton, Suyash Shringarpure, Chao Tian, Joyce Y. Tung, Vladimir Vacic, and Catherine H. Wilson. This work was supported by the National Human Genome Research Institute of the National Institutes of Health (grant number R44HG006981). The iPSYCH (The Lundbeck Foundation Initiative for Integrative Psychiatric Research) team acknowledges funding from The Lundbeck Foundation (grant no R102-A9118 and R155-2014-1724), the Stanley Medical Research Institute, the European Research Council (project no: 294838), the Novo Nordisk Foundation for supporting the Danish National Biobank resource, and grants from Aarhus and Copenhagen Universities and University Hospitals, including support to the iSEQ Center, the GenomeDK HPC facility, and the CIRRAU Center. A full list of the authors and affiliations in the iPSYCH-Broad autism group is provided in the Supplementary Information.

